# Extensive transgressive gene expression in testis but not ovary in the homoploid hybrid Italian sparrow

**DOI:** 10.1101/2021.04.15.440043

**Authors:** Homa Papoli Yazdi, Mark Ravinet, Melissah Rowe, Glenn-Peter Sætre, Caroline Øien Guldvog, Fabrice Eroukhmanoff, Alfonso Marzal, Sergio Magallanes, Anna Runemark

## Abstract

Hybridization can result in novel allelic combinations which can impact the hybrid phenotype through changes in gene expression. While mis-expression in F1 hybrids is well documented, how gene expression evolves in stabilized hybrid taxa remains an open question. As gene expression evolves in a stabilizing manner, break-up of co-evolved *cis*- and *trans*-regulatory elements could lead to transgressive patterns of gene expression in hybrids. Here, we address to what extent gonad gene expression has evolved in an established and stable homoploid hybrid, the Italian sparrow (*Passer italiae*). Through comparison of gene expression in gonads from individuals of the two parental species (i.e., house and Spanish sparrow) to that of Italian sparrows, we find evidence for strongly transgressive expression in male Italian sparrows - 22% of testis genes exhibit expression patterns outside the range of both parent species. In contrast, Italian sparrow ovary expression was similar to that of one of the parent species, the house sparrow (*Passer domesticus*). Moreover, the Italian sparrow testis transcriptome is 26 times as diverged from those of the parent species as the parental transcriptomes are from each other, despite being genetically intermediate. This highlights the potential for regulation of gene expression to produce novel variation following hybridization. Genes involved in mitochondrial respiratory chain complexes and protein synthesis are enriched in the subset that is over-dominantly expressed in Italian sparrow testis, suggesting that selection on key functions have molded the hybrid Italian sparrow transcriptome.

## Introduction

Hybridization between recently diverged species is an important evolutionary phenomenon resulting in novel gene combinations that become available to selection (Abbott et al., 2013; Mallet, 2005; Runemark, Vallejo-Marin, & Meier, 2019; Taylor & Larson, 2019). While bringing together diverged parental alleles in hybrids can lead to reduced hybrid fitness (Dobzhansky, 1936; Muller, 1942), it is now clear that hybridization is frequent and contributes to novel adaptive phenotypes (Marques, Meier, & Seehausen, 2019; Runemark et al., 2019; Taylor & Larson, 2019). Despite the growing body of evidence documenting admixed genomes, it is not clear how such intermediate genomes give rise to transgressive hybrid phenotypes (Rieseberg, Archer, & Wayne, 1999). Gene expression is the key intermediate step between genotype and phenotype, and divergence in regulation of gene expression is thought to play an important role in phenotypic evolution (Hodgins-Davis, Rice, & Townsend, 2015; Rest et al., 2013). Despite this, we lack an understanding of how gene expression evolves and contributes to novel phenotypes in hybrid taxa.

Gene expression evolves in a stabilizing manner, where regulatory elements accumulate mutations that keep gene expression at an optimum level for physiological functions (Coolon, McManus, Stevenson, Graveley, & Wittkopp, 2014; Gilad, Oshlack, & Rifkin, 2006; Hodgins-Davis et al., 2015). Specifically, changes in distal, *trans*-regulatory elements are compensated by changes in proximate, *cis*-regulatory elements (Landry et al., 2005). Such compensatory evolution means that as hybridization breaks up co-inheritance of regulatory elements, hybrids may experience novel combinations (Landry et al., 2005; Renaut, Nolte, & Bernatchez, 2009). The novel combinations of regulatory elements arising during hybridization has the potential to lead to transgressive gene expression, that transcends the range of parental expression profiles.

Hybrids resulting from strongly divergent taxa may exhibit extensive levels of transgressive expression arising from uncoupling of co-evolved *cis-* and *trans-*regulatory elements in F1 hybrids (Coolon et al., 2014; Haerty & Singh, 2006; McManus et al., 2010). In post-F1 generation hybrids, transcriptomes show a higher level of transgressive expression due to uncoupling of regulatory elements in the process of recombination. In the lake white fish *Coregonus clupeaformis* for example, F2 hybrids showed a higher level of nonadditive inheritance in gene expression compared to F1 hybrids (Renaut et al., 2009). An additional line of evidence of a non-linear relationship between genetic composition and gene expression comes from recent studies on allopolyploid plant taxa, hybrid taxa with doubled chromosome number. Both preferential expression of one of the parental genomes (Edger et al., 2017), and tissue dependent parental dominance in expression pattern (Kryvokhyzha et al., 2019) have been documented in allopolyploids. Jointly, the transgressive expression in experimental hybrids and non-linear relationship between genomic composition and gene expression similarity, suggest that gene expression will not be proportional to the parental contributions to the genome. Whether transgressive patterns of gene expression are found in homoploid hybrid taxa, hybrids without increase in ploidy, is unknown.

Here, we leverage a unique study system to investigate the nature of gene expression in a stabilized hybrid taxon, the homoploid hybrid Italian sparrow (*Passer italiae*), where thousands of generations of selection have resulted in a stable genome composition. The Italian sparrow arose from hybridization between the house sparrow (*Passer domesticus*) and Spanish sparrow (*Passer hispaniolensis*) (Fig. 1) (Elgvin et al., 2017; Hermansen et al., 2014; Hermansen et al., 2011; Trier, Hermansen, Saetre, & Bailey, 2014). The parent species diverged approximately 0.85 million years ago (Ravinet et al., 2018) and the Italian sparrow originated within *ca*. 5800 years, most likely as the house sparrow expanded its range into Europe and hybridized with the Spanish sparrow (Ravinet et al., 2018). This system provides a unique possibility to test to what extent a homoploid hybrid species has diverged in gene expression from the parent species after thousands of generations. In this study, we compared the gene expression profiles of the gonads of wild individuals of Italian sparrow to gene expression of the parental species to address if gene expression in Italian sparrows is intermediate to that of its parent species. At the same time, we opportunistically compared gene expression profiles to experimental F1 hybrid crosses (house x Spanish) bred in captivity. Thereby, we test whether gene expression reflects the intermediate genome composition of the hybrid Italian sparrow, or if it is transgressive for specific gene categories.

**Figure 1.**
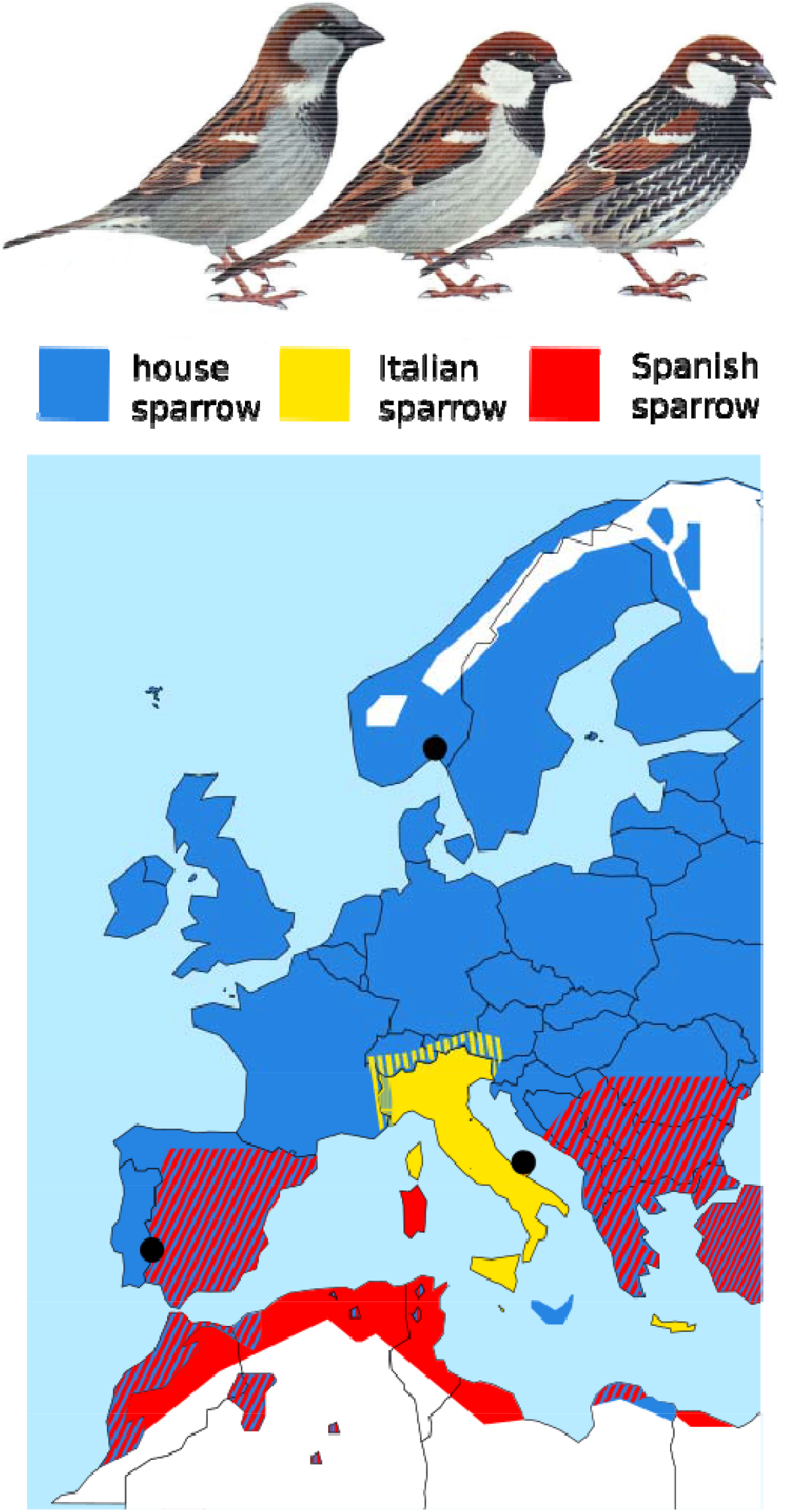
Species distribution and sampling locations. Top: Illustrations of male plumage patterns in house, Italian, and Spanish sparrows modified from (Svensson, Grant, Mullarney, & Zetterstroem, 1999). Bottom: A distribution map of house, Italian, and Spanish sparrows throughout Europe and northern Africa (Summers-Smith, 1988). This picture is modified from (Elgvin et al., 2017).

## Materials and methods

### Sampling, RNA extraction, library preparation, and sequencing

We compared the gene expression of wild Italian sparrows to wild individuals of its two parent species, the house and Spanish sparrow. Although the species differ in their distribution and phenology, all birds were sampled during the breeding season when birds were actively breeding (Supplementary section 1) at their respective locations. Spanish sparrows were sampled near Olivenza, Spain (38º 40’ 56”N, 7º 11’ 17”W) in March 2016, house sparrows in Oslo, Norway (59º 55’ 2”N, 10º 46’ 11”E) in May 2016 and Italian sparrows from Masseria Montanari in the Gargano peninsula (41º 54’ 36.8”N, 15º 51’ 13.0”E) in May 2016 (Fig. 1). All birds were trapped using mist nets and placed in a soft cotton bird bag until processing. All birds were euthanized by cervical dislocation, and gonads (ovary and the left testis) were harvested immediately after death. Sampling was performed between 10:00 and 16:00, with individuals from each of the species being sampled over most of the day, in order to avoid biases in sampling time. The gonads were stored in RNAlater, and immediately stored at -80 degrees, with extractions later performed at the University of Oslo using the Qiagen miRNeasy micro kit. Samples were prepped with TruSeq® Stranded mRNA library prep from Illumina on an automated Perkin Elmer Sciclone NGSx liquid handler and sequenced in the Norwegian Sequencing Center in Oslo, as paired-end (2×150 bp) on the Illumina Hiseq4000. The number of reads obtained per each sample is reported in Table S1.

### Read mapping, quality check and counting

We mapped reads from samples of all three groups (testis sample size: house sparrow = 5, Spanish sparrow = 5, Italian sparrow = 5; ovary sample size: house sparrow = 5, Spanish sparrow = 3 and Italian sparrow = 5) to the house sparrow reference genome using STAR2 v2.7.2b (Dobin et al., 2013) with default parameters and the house sparrow gene annotation as reported in (Elgvin et al., 2017). Over 90% of the reads mapped successfully (Table S1). The house sparrow general feature format (GFF) file contained 14,734 annotated genes of which 92.52% were anchored to chromosomes and 7.48% were located on unanchored scaffolds. Of the genes anchored to chromosomes, 12,595 genes were annotated on autosomes and 598 genes were on Z chromosome (Table S2). To ensure homogeneity in quality across our samples, we used RSeQC (Wang, Wang, & Li, 2012) to measure RNA integrity at the transcriptome level for each sample by calculating Transcript Integrity Number (TIN). The median TIN scores across samples ranged between 73.7 and 85.9 (Table S3). We counted the number of reads mapping to the house sparrow gene features using HTSeq v0.9.1 (Anders, Pyl, & Huber, 2015). Read counting was performed for all reads with minimum quality score of 30 and configured to handle reverse-stranded sequencing data, with the parameter controlling for overlapping gene features set to union. We created a dendrogram to show sample relationships. The main split was clearly between tissue types, i.e., testis and ovary (Fig. S1).

### Differential gene expression analysis

We analyzed the raw read counts obtained from htseq-count using the R package DESeq2 (Anders & Huber, 2010; Love, Huber, & Anders, 2014). We first tested genes for differential expression between house and Spanish samples. We then compared each of the parents to the Italian sparrow. We pre-filtered data to a minimum of total read counts of 10 across at least half of the samples in each pairwise comparison. To generate more accurate estimates of log2 fold change (LFC) for genes with low count number or large dispersion, we shrank LFC estimates. We considered genes to be differentially expressed if they showed False Discovery Rate (FDR) Padj < 0.05 and shrunken LFC > 0.32. All statistical analyses were completed using R version 4.0.2 (R Core Team 2020).

### Classification of gene expression inheritance

The mode of inheritance for differentially expressed genes was determined following (McManus et al., 2010). We normalized gene expression using median of ratios method in DESeq2 (Love et al., 2014). Genes whose total expression in Italian sparrow deviated significantly more than 1.25-fold (LFC > 0.32) from that of either parent were considered to have non-conserved inheritance. These genes were classified as having additive, dominant, under-dominant, or over-dominant inheritance, based on the magnitude of the difference between total expression in the Italian sparrow and in each parental species. Genes for which expression in the Italian sparrow was less than for house sparrows and greater than for Spanish sparrow (or vice versa) were classified as additive; genes for which expression in Italian sparrow was similar to one of the parents were classified as dominant; and genes for which expression in Italian sparrow was either greater than or less than both parent species were classified as over-dominant and under-dominant (transgressive), respectively (Figs. 3A-3B).

### Functional Annotation and Enrichment Analysis

Gene ontology (GO) analysis was done for differentially expressed genes in each pairwise comparison. GO functional annotations and gene descriptions were obtained for protein sequences from the entire house sparrow protein set as described in (Rowe et al., 2020) using PANNZER (Koskinen, Toronen, Nokso-Koivisto, & Holm, 2015). Functional enrichment of GO terms present in the set of differentially expressed genes relative to the background consisting of all genes expressed and tested for differential expression was performed using the clusterProfiler (Yu, Wang, Han, & He, 2012) and significant enrichment was determined at FDR < 0.05. Interactions among proteins with significant GO terms were predicted using STRING-v11 (Szklarczyk et al., 2019) using “Experiment”, “Databases” and “Co-expression” as interaction sources. We set the minimum required interaction score to the highest confidence (0.9) and used only the query proteins to build the interaction network after removing the disconnected nodes in the network. The obtained network was then exported to Cytoscape (Shannon et al., 2003) and major representations of biological processes were detected using ClueGO (Bindea et al., 2009) with Padj< 0.01.

### Analysis of experimental hybrids

Although our study was designed to allow the comparison of wild Italian sparrows to wild individuals of its two parent species, the house and Spanish sparrow, we also capitalized on the availability of captive-bred experimental F1 hybrids produced by crossing house sparrow females and Spanish sparrow males (Supplementary section 2, Fig. S2). As experimental hybrids were bred in an area where house and Spanish sparrows are sympatric for at least part of the year (Olivenza, Spain; see Fig. 1) and correct identification of females can be challenging, we performed a post-sampling genomic analysis of all samples (Supplementary section 3). The mtDNA of experimental F1 hybrids were correctly grouped with house sparrow (Fig. S3). However, the F1 hybrids, did not show the expected equal ancestry ratio from each parental species and the PCA with the Z chromosome of F1 females showed clustering with Z chromosome from house sparrow (Figs. S4-S5). We therefore acknowledge that the comparison between these experimental hybrids and the other species sampled in our study is suboptimal. Nonetheless, since the comparison of the stabilized homoploid hybrid Italian sparrow and artificial early generation hybrids between house and Spanish sparrows provides an interesting contrast, we decided to retain the experimental hybrid samples while clarifying the limitations to how these findings can be interpreted. All gene expression analyses were carried out in these experimental hybrids in same way as described for the Italian sparrow samples.

## Results

### Differential expression between the parental species

Gene expression divergence in gonads between the parental species was strongly asymmetric with testis showing a more conserved pattern of expression compared to ovary (Table 1). In the testis, 135 genes (1.16%) and in ovary, 1382 genes (11.65%) were significantly differentially expressed. Of the differentially expressed genes, 60.7% of genes in testis and 69.3% of genes in ovary were upregulated in the Spanish sparrow. A significantly higher proportion of genes that were differentially expressed in testis were located on the Z chromosome (24 genes, hypergeometric test, *P*-value < 1.82-08). We found no evidence for functional enrichment among differentially expressed genes in parental species for either testis or ovary.

**Table 1.** Number of differentially expressed genes and log2 fold change (LFC) with Padj< 0.05^**†**^ and LFC^‡^ > 0.32.

| Comparison | Testis |  |  | Ovary |  |  |
| --- | --- | --- | --- | --- | --- | --- |
|  | Significant | LFC > 0 | LFC < 0 | Significant | LFC > 0 | LFC < 0 |
| Spanish - house | 135 | 82 | 53 | 1382 | 958 | 424 |
| Italian - house | 3536 | 1962 | 1574 | 22 | 13 | 9 |
| Italian - Spanish | 3581 | 1779 | 1802 | 1508 | 394 | 1114 |

$^\dagger$ Results with $\text{Padj} < 0.01$ are reported in Table S4
$^\ddagger$ LFC: Log Fold Change

### Differential expression in Italian sparrows compared to the parental species

In contrast to the intermediacy of the Italian sparrow genome, Italian sparrow testis gene expression differed more in comparison to the parental species than the parent species differed from each other (Table 1). Specifically, approximately one-third of the testis transcriptome was differentially expressed in Italian sparrow compared to both parental species, whereas ovary expression was similar to that of the house sparrow (Table 1, Fig. 2). In the Italian sparrow testis, 3536 genes (30.45%) and 3581 genes (30.9%) were differentially expressed in comparison to house and Spanish sparrow, respectively. For Italian sparrow ovary, only 22 genes (0.18%) differed from the house sparrow whereas 1508 genes (12.63%) were differentially expressed compared to the Spanish sparrow. Significant over-representation of Z-linked genes among the differentially expressed genes was detected in testis, but only in the comparison to Spanish sparrow (196 of the 3581 genes, hypergeometric test, *p* value = 0.006). Most genes in both testis and ovary were up-regulated compared to house and down-regulated compared to Spanish sparrow (Chi-squared test *p* value = 1.05e-06 in testis and 0.001 in ovary) (Fig. 2).

**Figure 2.**
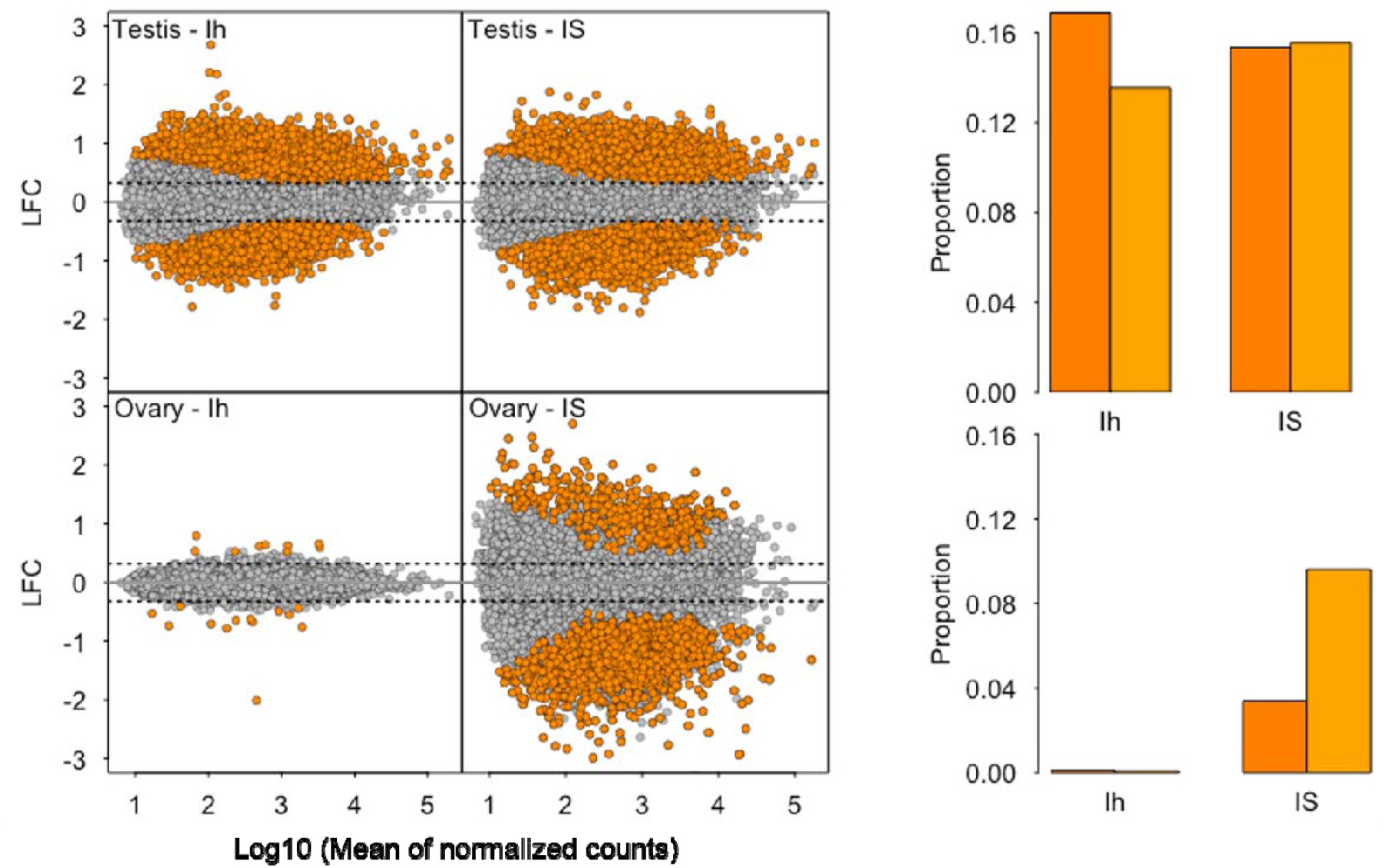
Gene expression in Italian sparrow in comparison to house (Ih) and Spanish (IS) for testis (top) and ovary (bottom). Log2 fold change (LFC) is plotted as a function of mean of normalized counts. Bar plots summarize the proportion of up- and down-regulated genes. Dark orange: up-regulated, light orange: down-regulated.

**Figure 3.**
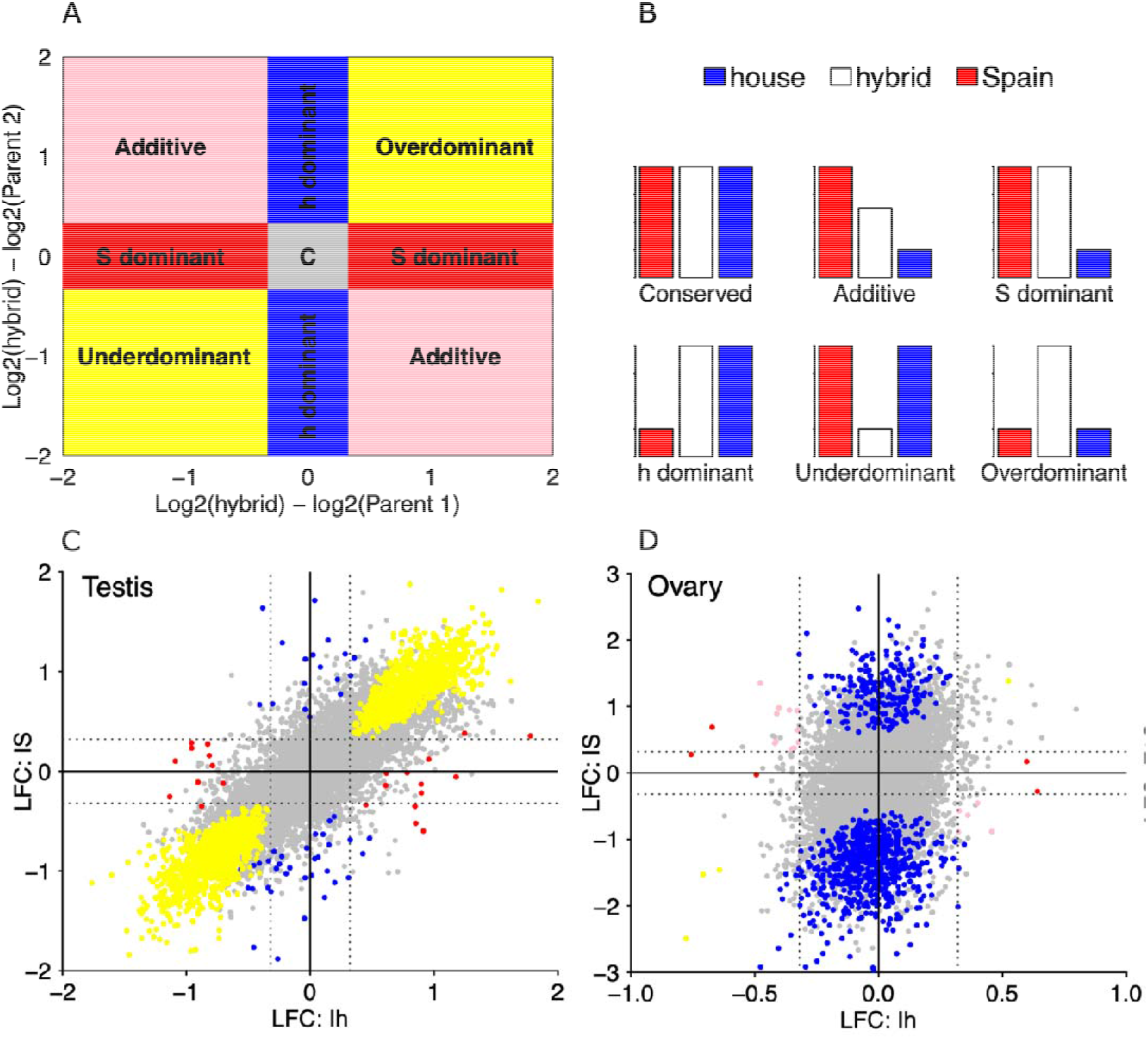
Inheritance pattern of gene expression in testis and ovary in Italian sparrows. A and B: Schematic figures representing the classification of inheritance patterns. C and D: Scatter plots showing shrunken log2 fold change (LFC) in testis between the Italian sparrow with the house sparrow (Ih) on the *x*-axis and with Spanish sparrow on the *y*-axis (IS), respectively. D: Scatter plots show shrunken log2 fold change (LFC) in ovary between the Italian sparrow with the house sparrow on the *x*-axis and with Spanish sparrow on the *y*-axis, respectively.Grey points in each graph depicts the total number of genes studied for gene expression with those colored representing the ones significantly different from parental species to be divided into each of the inheritance categories (Conserved: grey, additive: pink, house dominant: blue, Spanish dominant: red, transgressive (over-dominant and under-dominant: yellow). Grey dotted lines indicate the log2 fold-change threshold of 0.32 used for classification of inheritance pattern.

Genes that were differentially expressed in Italian sparrow testis were enriched for similar biological processes in comparison to both parent species. Primarily, functions involving protein synthesis and mitochondrial gene expression and function were overrepresented. Comparisons between the Italian sparrow and the house sparrow yielded 27 significant GO categories (Table S5), with top biological processes involved in viral transcription, SRP-dependent co-translational protein targeting to membrane, translation, ribosome and mitochondrial respiratory chain complex I assembly. In comparisons between the Italian sparrow and the Spanish sparrow, the 15 enriched GO terms reflected similar biological processes, with ribosome as the top GO term (Table S6). In ovary, 6 GO terms were detected in the comparison to house sparrow with sodium ion transmembrane transporter activity as the top term (Table S7) and 1 GO term, negative regulation of protein phosphorylation detected in the comparison to Spanish sparrow (Table S8).

### Classification of inheritance patterns of gene expression

Gene expression was generally conserved in Italian sparrow (Table 2), but in the testis of Italian sparrow, 2611 genes (22.71%) showed a non-conserved pattern of inheritance with transgressive expression and house-dominant being the two largest categories (Fig. 3C). Transgressively expressed genes in Italian sparrow comprised 22% of the testis transcriptome tested for inheritance. Over 96% of genes with non-conserved pattern of inheritance in Italian sparrow ovary, had a house-dominant pattern of expression (Fig. 3D). In contrast to the high incidence of transgressive expression observed in Italian sparrow testis, only 3 genes (0.028%) were transgressively expressed in ovaries. Over-dominant genes in Italian sparrow testis were enriched for functional categories involved in protein synthesis, mitochondrial protein complex and gene expression and binding of sperm to zona pellucida (Table S10). We found a gene network with significant protein-protein interaction (PPI Enrichment: 1.0E-16) among the over-dominant genes in Italian sparrow (Fig. 4). A different set of genes was under-dominant in Italian sparrow, with functions including regulation of cellular component size and negative regulation of neuron projection development.

**Table 2.** Number and percentage of genes in each inheritance category^**†**^.

| Inheritance | Testis | Ovary |
| --- | --- | --- |
|  | Italy | Italy |
| Conserved | 8883 (77.30%) | 10889 (92.26%) |
| Additive | 2 (0.02%) | 19 (0.16%) |
| house dominant | 54 (0.47%) | 895 (7.58%) |
| Spanish dominant | 25 (0.22%) | 7 (0.06%) |
| Under-dominant | 1205 (10.47%) | 3 (0.02%) |
| Over-dominant | 1325 (11.53%) | 1 (0.008%) |

$^\dagger$ Inheritance classification for differentially expressed genes with $\text{Padj} < 0.01$ is reported in Table S9

**Figure 4.**
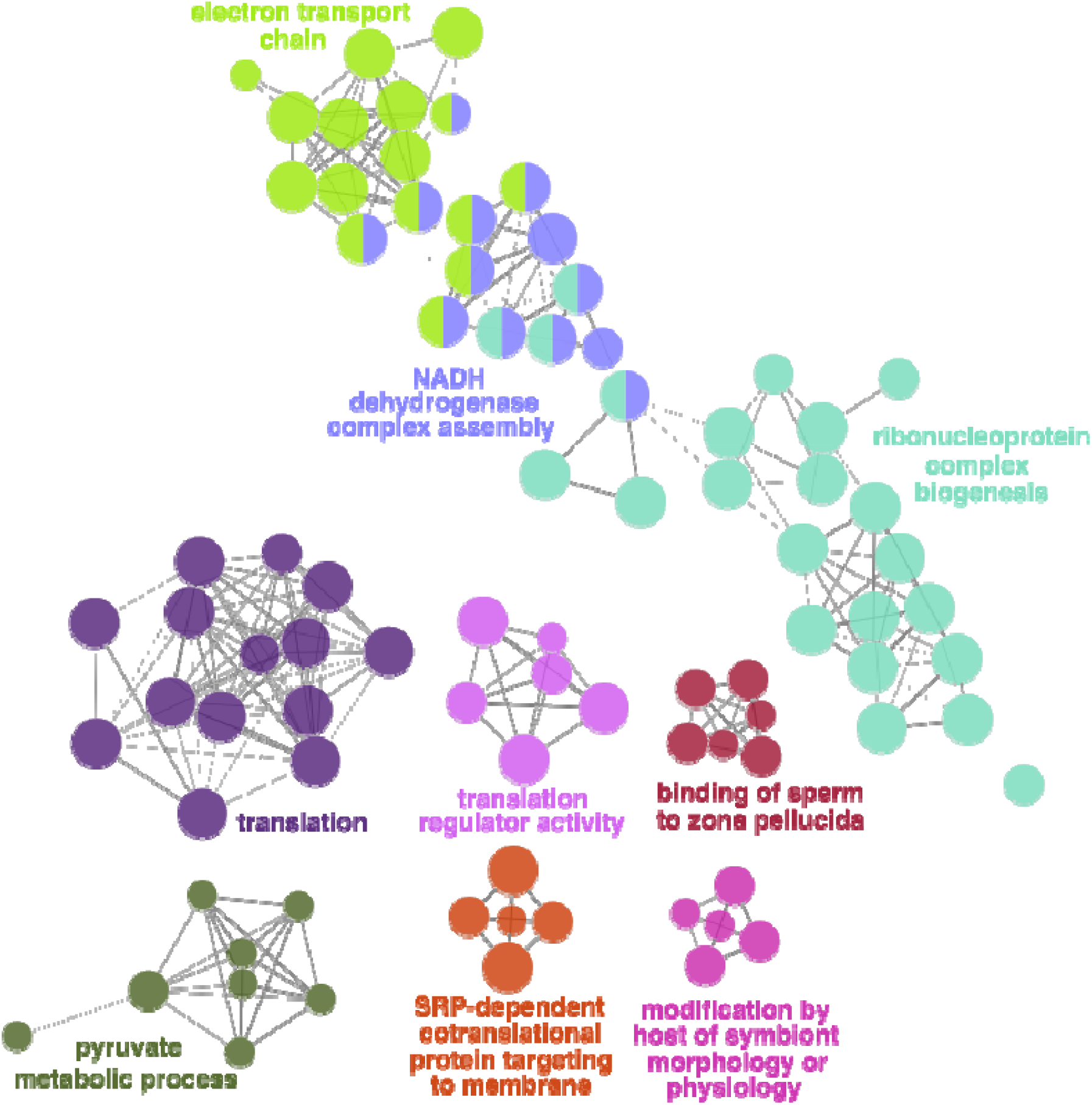
Gene ontology network represented in over-dominant genes in testis of Italian sparrow. Different colors depicts different significant biological processes. Circle size represents adjusted *P*-value for each node with all Padj < 0.01 where node size represents the term enrichment significance.

### Patterns of expression in experimental hybrids

The overall magnitude of differences in gene expression in the experimental hybrid compared to both parent species was much lower than for the Italian sparrow (Figure S6). In testis, 106 genes (0.9%) and 263 genes (2.25%) were differentially expressed compared to house and Spanish sparrows, respectively. In ovary, 30 genes (0.25%) and 140 genes (1.15%) showed a significant difference in expression compared to the house and Spanish sparrows, respectively. Moreover, in the experimental hybrid testis only 0.37% of the genes were transgressively expressed (Table S11). Gene expression in experimental hybrid ovary also was more conserved in relation to the parent species compared to that of the Italian sparrow, with the two largest categories of inheritance being additive and house-dominant (Table S11). Additional results from the experimental hybrids are presented in the Supplementary section 4 and Tables S12-S15.

## Discussion

This study presents, to the best of our knowledge, the first evidence for transgressive gene expression in a wild homoploid hybrid species. The strongly divergent gene expression profile in Italian sparrow testis highlights the potential of intermediate hybrid genomes to rapidly evolve through altered regulation of gene expression. About 22% of testis transcriptome in Italian sparrow showed transgressive expression pattern, while this level was less than 0.03% in the ovaries. Transgressive gene expression, despite conserved parental expression, could arise in later generation hybrids due to the uncoupling of co-evolved *cis*- and *trans*-regulatory elements through recombination (Landry et al., 2005; Signor & Nuzhdin, 2018; Takahasi, Matsuo, & Takano-Shimizu-Kouno, 2011). There are, however, few studies of gene expression in post F1-hybrids and transgressive expression is commonly referred to “mis-expression” in early generation hybrids (Signor & Nuzhdin, 2018). Here, we show that the homoploid hybrid Italian sparrow has achieved a 26 times stronger divergence in testis gene expression during ca. 5800 years of evolution following hybridization than the parents have accumulated during 0.68 million years (Ravinet et al., 2018). This might imply that sorting of parental regulatory elements following hybridization can produce novel expression phenotypes from intermediate genomes.

Hybrid incompatibilities leading to reproductive isolation due to novel, untested genetic combinations are most commonly thought to occur when parental species are fixed for different alleles (Cutter, 2012). However, transgressive expression patterns in the hybrids of more recently diverged species pairs can also arise either due to polymorphic incompatible loci (Cutter, 2012) or physiological response to hybrid dysfunction (Barreto, Pereira, & Burton, 2015). In interpopulation hybrids at F2 or later generation of copepod species (*Tigriopus californicus*), transgressive gene expression mainly reflected physiological responses in hybrids (Barreto et al., 2015). Therefore, given the relatively low divergence between house- and Spanish sparrow, transgressive expression in the Italian sparrow could also be due to bringing together polymorphic *cis*- and *trans*-regulatory variants from the parental species. Alternatively, transgressive gene expression may arise from lineage specific mutations that have been favored by selection in Italian sparrow. Each of these hypotheses can be evaluated in future work by studying the optimal F1 and F2 hybrids and their parental individuals to identify *cis* and *trans*-regulatory variants.

The degree of difference in gene expression was parent- and tissue specific. We generally observed fewer differentially expressed genes relative to house sparrow than relative to the Spanish sparrow for the Italian sparrow, and this was especially true for ovarian tissue. While smaller sample size in Spanish sparrow might have contributed to the house-dominant pattern of expression, it is likely that our findings are at least partly explained by the fact that the Italian sparrow has house sparrow mitochondria (Hermansen et al., 2014; Runemark, Eroukhmanoff, Nava-Bolanos, Hermansen, & Meier, 2018; Trier et al., 2014). Mito-nuclear interactions can strongly affect gene expression (Sanchez-Ramirez, Weiss, Thomas, & Cutter, 2021), and even result in mis-expression (Runemark, Eroukhmanoff, et al., 2018). Mito-nuclear genes have a role in reproductive isolation in Italian sparrows and have been under selection for inheritance of the house sparrow allele (Hermansen et al., 2014; Runemark, Trier, et al., 2018; Trier et al., 2014). Interestingly, we found over-expression of genes involved in functions related to mitochondrial respiratory chain, cytosolic and mitochondrial ribosomal proteins in the hybrid Italian sparrow. Correct mitochondrial function requires a coordinated expression of both nuclear and mitochondrial genomes (Ryan & Hoogenraad, 2007). Therefore, the upregulation of genes involved in respiratory chains and protein synthesis corroborate the evidence for a strong effect of hybridization on genes associated with metabolism in other bird taxa (Wagner, Curry, Chen, Lovette, & Taylor, 2020). Overexpression of cytosolic ribonucleoproteins and mitochondrial respiratory chain enzymes has also been observed in later generation hybrids (F3+) in copepod species and has been linked to cellular responses to physiological dysfunction (Barreto et al., 2015). Our study thus contributes to the growing body of evidence of mito-nuclear interactions as important reproductive barriers (Hill, 2017), and provides a unique insight into altered patterns of gene expression as a possible resolution to the conflict.

In addition to mito-nuclear genes, over-dominant genes in Italian sparrow contained ribonucleoproteins present in both cytosolic and mitochondrial ribosomes. RNA binding proteins play an important role in post-translational regulation of gene expression and spermatogenesis and appear to be key regulatory factors that ensure male fertility (Paronetto & Sette, 2010; Phillips et al., 2019). Several of the over-dominant genes also coded for subunits of T-complex protein Ring Complex (TRiC) involved in folding of about 10% of the proteome. Subunits of TRiC complex are required for spermatogenesis (Counts, Hester, & Rouhana, 2017) and are under positive selection among the seminal fluid genes in passerine species (Rowe et al., 2020).

Although we suggest caution in interpretation of our experimental F1 hybrid data due to the unusual genomic composition of F1 hybrids, our work is in agreement with findings from F1s in two other songbird species (Davidson & Balakrishnan, 2016; Mugal et al., 2020). F1 hybrids of flycatchers also showed a tissue specific pattern of expression differences, with a low degree of transgression in genes expressed in testis (Mugal et al., 2020), similar to the pattern in our experimental hybrids but contrasting to the strong transgression in the testis transcriptome of the Italian sparrow. Future studies should therefore extend sampling to non-reproductive tissues in order to assess whether transgressive expression is consistent across tissues. While a standardized environment and time for sampling would be ideal, this is not easily achieved in wild species that differ in geographic distributions, phenologies and thermal comfort zones, as sampling at the same time could capture different phenological stages. Future studies should sample these species over time to remedy this complication.

Homoploid hybrid species are expected to have higher level of genetic variance due to homologous recombination leading to transgressive phenotypes (Mallet, 2007; Rieseberg et al., 2003). The extensive level of transgressive expression in the Italian sparrow testis is therefore in agreement with this expectation. Gene expression is the key intermediate step between genotype and phenotype, therefore examining the link between transgressive gene expression and transgressive hybrid phenotypes is a crucial next step which will significantly increase our understanding of how novel variation arises from hybridization and the role of gene expression divergence in the evolution of reproductive isolation from parental species.

## Acknowledgements

We thank Angelica Cuevas, Matteo Caldarella, Severino Vitulano and Giuseppe Cardone for assistance with sampling, which was done in collaboration with the Puglia region, Centre for Naturalistic Studies ONLUS (CSN, Italy) and ISPRA (Italy). We are grateful to Richard Bailey and Daniel Hooper for their comments on earlier versions of this manuscript. The computations were enabled by resources in project SNIC2020-6-222 provided by the Swedish National Infrastructure for Computing (SNIC) at UPPMAX, partially funded by the Swedish Research Council through grant agreement no. 2018-05973. This work was financed by a Norwegian Research council grant to G-P. S. and A. R. and a Wenner-Gren and a Swedish Research Council grant to A.R.

## Data accessibility and benefit-sharing statement

Raw RNA sequencing reads will be deposited in the SRA (BioProject XXX) upon acceptance of the manuscript for publication. R script containing code for differential gene expression analyses and the figures presented in this manuscript is deposited under github.com/Homap/Expression_sparrow. In this work, all samples taken from Italy and Spain to Oslo in accordance with the Nagoya protocols. In Spain, all trapping and sampling of birds was conducted in accordance with Spanish Animal Protection Regulation RD53/2013, and all methods were approved by the Institutional Commission of Bioethics at the University of Extremadura (CBUE 49/2011). Birds were sampled in Oslo with permission from Miljødirektoratet (2016/2225). In Italy, all birds were sampled with permission from ISPRA (protocol number 12404) and with permission (no. 305) from Puglia region. Research was performed in collaboration and cooperation with local researchers from University of Extremadura. All collaborators are included as co-authors. International collaboration has been fundamental to this work and has contributed to sharing of scientific results, cooperation, education, and training.

## Author contributions

A.R. and H.P.Y. conceived of the study. A.M. and S.M. raised the experimental F1 hybrids and M. Ro., M. Ra., A.R., F.E, C.O.G and G-P. S. performed field work and sampling, C.O.G performed the molecular lab work, and H.P.Y. designed and performed the bioinformatical analysis based on discussions with A.R, M. Ra., and M.Ro. H.P.Y. drafted the manuscript, with revisions based on the input from all co-authors.

## Notes

### Competing Interest Statement

The authors have declared no competing interest.

### Summary of Updates

In this version, we have added more detailed analyses and description about the sampling strategy and experimental F1 hybrid ancestry and have worked further on providing explanations for our results.

