## Supplementary Materials for "Extensive transgressive gene expression in testis but not ovary in the homoploid hybrid Italian sparrow"


| | |
|---|---|
| Supplementary section 1: Sampling of reproductive tissues and timing of reproduction in sparrows | 2 |
| Supplementary section 2: Breeding and sampling of experimental F1 hybrids | 3 |
| Supplementary section 3: Genetic characterization of samples | 3 |
| Supplementary section 4: Gene expression pattern and its mode of inheritance in the experimental F1 hybrids | 4 |
| Supplementary References | 5 |
| Table S1: Summary of RNA-seq read statistics | 6 |
| Table S2: Number of annotated genes per chromosome of house sparrow | 8 |
| Table S3: Transcript Integrity Number (TIN) | 9 |
| Table S4: Number of differentially expressed genes and log2 fold change (LFC), padj<0.01 | 10 |
| Table S5: Significant GO categories for the comparison Italian – house testis | 11 |
| Table S6: Significant GO categories for the comparison Italian – Spanish testis | 13 |
| Table S7: Significant GO categories for the comparison Italian – house ovary | 14 |
| Table S8: Significant GO categories for the comparison Italian – Spanish ovary | 15 |
| Table S9: Number and percentage of genes in each inheritance category for genes in Table S4 | 16 |
| Table S10: Significant GO categories for over-dominant genes in testis of Italian sparrow | 17 |
| Table S11: Number and percentage of genes in each inheritance category in experimental F1 hybrids | 19 |
| Table S12: Significant GO categories for the comparison of experimental F1 – house testis | 20 |
| Table S13: Significant GO categories for the comparison of experimental F1 – Spanish testis | 22 |
| Table S14: Significant GO categories for the comparison experimental F1 – Spanish ovary | 23 |
| Table S15: Significant GO categories for over-dominant genes in testis of experimental F1 hybrid | 25 |
| Figure S1: Cluster dendrogram of testis and ovary samples | 26 |
| Figure S2: Pictures of two of experimental F1 hybrid males | 27 |
| Figure S3: Mitochondrial tree | 28 |
| Figure S4: Principal component analysis of autosomal SNPs | 29 |
| Figure S5: PCA of the ovary Z chromosome | 30 |
| Figure S6: Gene expression and inheritance pattern in experimental F1 hybrids | 31 |

**Supplementary section 1**

**Sampling of reproductive tissues and timing of reproduction in sparrows**

We sampled adult house, Italian and Spanish sparrows at a range of sites across Europe. Although the species were sampled at different time periods, all birds were sampled during the breeding season for their own population. Active breeding was confirmed in all birds through examination of reproductive organs and tissues and observation of the local population for breeding activity (e.g., copulation, nest building, fledglings, etc). House sparrows were sampled in May 2016 in Oslo. All sampled birds had enlarged gonads and males were actively producing sperm. Broadly speaking, temperate zone populations of the house sparrow exhibit a prolonged breeding season lasting from March through to early September (Anderson 2006). House sparrows are multi-brooded and highly asynchronous breeders. Thus, within a population breeding pairs will be at different reproductive stages. (e.g., nesting, incubating, raising nestlings, fledglings). For males, testis size increases rapidly in February/March and testes remain large until late in the breeding season (*c.* August; Anderson 2006). The population of house sparrows used in this study regularly breeds from April to late July, and in some years into late August (M. Rowe, personal observation). Males can be actively producing sperm as early as late March (M. Rowe, unpublished data). During sampling for this project, the population was observed to be in peak breeding activity, with a range of breeding stages observed (e.g., nest building, copulations).

Italian sparrows were sampled in May 2016 in Italy. All sampled birds had enlarged gonads at the time of sampling and were actively producing sperm. Average testis dimensions of sampled birds were (1) left testis: length 9.96 mms, width 7.59 mm; (2) right testis: length. 9.34 mm, width 7.87 mm. Detailed observations of breeding activity in a nearby population (Lago Salso) confirmed that Italian sparrows in this region were actively breeding, with a range of breeding stages observed (e.g., nest building, incubation, nestling care). Further, across Italy, populations of Italian sparrow have been observed breeding through August. Thus, the Italian sparrow also appears to have a prolonged breeding season and breeding is highly asynchronous.

Spanish sparrows were sampled in late March 2016 in Spain. As with the house sparrow, Spanish sparrows may raise multiple broods in one year. In Europe, the breeding season for Spanish sparrows is prolonged, running from March to August (Summers-Smith 1988). The population of Spanish sparrows used in this study showed clear signs of breeding activity (A. Marzal, personal observation). Birds captured had a hatching plate and the males had a more prominent cloaca, clear signs of being in the breeding season. In addition, nest building starts at the beginning of March and by the end of March many nests are already containing eggs. Importantly, all sampled birds had enlarged gonads. Although we lack measures of testis size for the birds used in this study, males collected in the subsequent year on the 28$^{th}$ of March (2017) had an average combined testes mass of 0.576 ± 0.11 grams (mean ± SD, range 0.478 – 0.688 grams). This is comparable to samples collected from Spanish sparrows at another sampling event (samples not used in the current study) during May; average combined testes mass of 0.625 ± 0.07 grams (mean ± SD, range 0.506 – 0.700 grams; M. Rowe unpublished data). Importantly, while we acknowledge that Spanish sparrows were sampled earlier in the breeding season than either the house or Italian sparrow, we are confident that all birds were actively breeding. Nonetheless, we consider the potential consequences of these different sampling times in our discussion in the main manuscript.

**Supplementary section 2**

**Breeding and sampling of experimental F1 hybrids**

Experimental F1 hybrids were bred in captivity using wild-caught house sparrow females and Spanish sparrow males. Birds were captured between 10th of January and 27th of February 2014. House sparrow females were captured at a farm approximately 10 km southeast of Olivenza, Spain (38° 38' 02"N, 7° 02' 59"W) early in the sampling period and generally prior to the arrival of Spanish sparrows migrating into the broader region. Unlike the house sparrow, the Spanish sparrow is known to migrate, though the extent of migration is highly variable across populations (Summers-Smith 1988). Importantly, Spanish sparrows have never been observed at this site or in the surrounding farms (S. Magallanes, personal observation). Sparrows have been captured in this area from 2014 to 2021 for different studies and no other species of sparrow than the house sparrow has ever been captured. Male Spanish sparrows were captured at a second farm site approximately 15km west of Olivenza (38° 38' 59"N, 7° 12' 59"W) after their migration into the region. Although Spanish and house sparrows co-occur at this site during the Spring and breeding season, males of the two species can be easily distinguished based on their plumage. Birds were placed in large outdoor aviaries in mixed-sex groups and provided with food and water *ad lib*. No breeding activity was observed during the first year of captivity, as seems to be common for wild-caught sparrows held in captivity (M. Rowe, personal observation). In 2015, captive birds bred successfully, producing several clutches of experimental F1 hybrid house sparrow (female) x Spanish sparrow (Male) offspring.

**Supplementary section 3**

**Genetic characterization of samples**

To genetically confirm the identity of all individuals, we conducted a phylogenetic analysis based on the mitochondrial RNA (mtRNA). We extracted mtRNA mapped reads to the house sparrow reference genome and used GATK HaplotypeCaller v4.1.4.1 (DePristo, et al. 2011) in the haploid ploidy mode to call variants for each individual sample. We kept only single nucleotide polymorphisms (SNPs) and removed SNPs that showed clustering of at least 3 SNPs in a window of 35 bases, SNPs with FS > 30, QD < 2.0, MQ < 40.0 and DP < 5. For each individual, we constructed mtDNA sequences by replacing SNPs on the mtDNA house sparrow reference with the called nonreference genotype using the consensus option in bcftools v1.10 (Li 2011). We aligned mtDNA sequences from all individuals and constructed a maximum likelihood phylogeny using the GTR model in Seaview (Gouy, et al. 2010). This analysis revealed that four of the experimental F1 samples cluster with Spanish sparrows and two Spanish sparrow samples cluster with house mtDNA (Fig. S3). To further investigate genome-wide patterns, we performed variant calling as described above for the nuclear genome, using the diploid ploidy mode. We genotyped the samples across testis and ovary separately and kept only biallelic sites with no missing information. This left us with 502,162 and 381,312 autosomal SNPs in testis and ovary, respectively. We used plink v1.90b4.9 (Purcell, et al. 2007) to prune SNPs for linkage disequilibrium by taking 100 Kb windows with a 20 SNP step size and pairwise $r^2$ threshold of 0.1 and conducted principal components analysis (PCA). This analysis confirmed the results from mtDNA tree with three F1 hybrid testis samples clustering with Spanish sparrow samples, two Spanish ovary samples clustering with F1 hybrid and one F1 hybrid ovary sample clustering with Spanish samples (Fig. S4). Italian sparrows and experimental F1 hybrids are expected to have house mtDNA while Spanish sparrows should have a separate cluster for their mtDNA. These samples were excluded from all downstream analyses, which left a sample size of testis: house sparrow: 5, Spanish sparrow: 5, Italian sparrow:5 and F1 hybrid: 5 and sample size of ovary: house sparrow: 5, Spanish sparrow: 3, Italian sparrow: 5, F1 hybrid: 8.

Experimental F1 hybrids were located more closely to the house sparrow than expected if the parental species were pure without any levels of introgression (Fig. S4). While the mtDNA analysis confirmed ancestry from house female consistent with experimental design, we found a surprising pattern for the Z chromosome. Since birds are female heterogametic with ZW chromosomes, females should receive their Z chromosome from their fathers. Since Spanish sparrow were the fathers in this design, we would expect the experimental F1 hybrids to cluster with them. However, we observed the clustering of female experimental F1 hybrids with the Z chromosome from house sparrows (Fig. S5). While the pictures captured of experimental hybrids show a clear crown coloration pattern similar to Spanish sparrow (brown crown in Spanish sparrow vs. the grey crown in house sparrow) (Fig. S2). Regardless of their complexity in their ancestry pattern, the data from the experimental F1 hybrids, although, suboptimal, provide an opportunity to compare level of expression divergence in early generation hybrid compared to a stabilized hybrid species, the Italian sparrow.

**Supplementary section 4**

**Gene expression pattern and its mode of inheritance in the experimental F1 hybrids**

We performed read mapping, differential gene expression, classification of inheritance patterns and analysis of gene ontology as described in the main text. Similar to the pattern observed in Italian sparrow, both testis and ovary showed a more conserved pattern of expression between the experimental hybrid and the house sparrow than between experimental hybrids and the Spanish sparrow (Fig. S6A). The overall magnitude of differences to the parent species was much lower than for the Italian sparrow. In testis, 106 genes (0.9%) and 263 genes (2.25%) were differentially expressed compared to house and Spanish sparrows, respectively. In ovary, 30 genes (0.25%) and 140 genes (1.15%) showed a significant difference in expression compared to the house and Spanish sparrows, respectively. Similar to the Italian sparrow, the pattern of up- and down-regulation was consistent in both testis and ovary in the comparison to each of the parental species. A larger number of genes were up-regulated in experimental hybrids in the comparison to house sparrow and down-regulated in the comparison to Spanish sparrow (Chi-squared test *P*-value = 1.96-05 in testis and 0.01 in ovary).

As in the Italian sparrow, there was an over-representation of Z-linked genes among those differentially expressed between experimental hybrids and Spanish sparrows in testis (31 of the 263 differentially expressed genes, hypergeometric test, *P*-value = 2.42-06). The enriched functions were different to those enriched for Italian sparrows. Testis genes differentially expressed in experimental hybrids showed functional enrichment for primarily collagen catabolic process. In the comparison to house sparrow, 46 GO terms were significantly over-represented. Among these GO terms, 20 GO terms had an Padj < 0.01, whereas there were few significant GO terms compared to Spanish sparrows with top GO term common with the comparison to house sparrow (Tables S12 and S13). The top 5 GO terms were enriched in proteinaceous extracellular matrix, extracellular matrix structural constituent, extracellular vesicle, collagen trimer and collagen catabolic process. In ovary, while there was no functionally significant term associated with differentially expressed genes in experimental hybrids compared to the house sparrow, there were 28 significant GO terms in the comparison to Spanish sparrow with the top category to be response to vitamin D (Table S14).

Similar to Italian sparrow, gene expression was generally conserved in experimental hybrids. In experimental hybrid testis only 0.37% of the genes were transgressively expressed (Fig. S6B). The majority of the genes with non-conserved pattern of inheritance in Italian sparrow ovary, over 96% of these genes, had a house-dominant pattern of expression. Gene expression in experimental hybrid ovary was more conserved compared to that of the Italian sparrow. Out of 376 genes with non-conserved pattern of inheritance, the two largest categories of inheritance were additive and house-dominant inheritance pattern. Over-dominant genes in experimental hybrid testis were functionally enriched for yet another set of GO-terms, including collagen catabolic process and proteinaceous extracellular matrix (Table S15). The only significant GO term for under-dominant genes in experimental hybrid testis was voltage-gated potassium channel complex.

**Table S1.** Summary of RNA-seq read statistics, total number of reads, percentage of uniquely mapped reads to the reference genome and percentage of reads mapped to genes (A) house sparrow (B) Spanish sparrow (C) Italian sparrow (D) experimental F1 hybrids.

(A)

| **Number of raw reads [$10^{06}$] / Uniquely mapped reads %/ Reads mapped to genes %** | | | |
|---|---|---|---|
| | Ovary | | Testis |
| **HTE10** | 36.340/ 92.77 / 57.015 | **HTE06** | 40.110/ 89.15/ 56.934 |
| **HTE02** | 37.803/ 92.70/ 56.834 | **HTE07** | 40.033/ 87.41/ 55.003 |
| **HTE03** | 34.491/ 93.33/ 55.419 | **HTE08** | 41.048/ 89.89/ 56.094 |
| **HTE04** | 34.002/ 93.40/ 56.704 | **HTE09** | 48.100/ 89.83/ 56.599 |
| **HTE05** | 38.228/ 93.60/ 56.695 | **HTE01** | 39.146/ 88.55/ 57.164 |
| **Mean** | 36.734/ 92.316/ 56.53 | **Mean** | 41.126/ 89.81/ 56.36 |

(B)

| **Number of raw reads [$10^{06}$] / Uniquely mapped reads %/ Reads mapped to genes %** | | | |
|---|---|---|---|
| | Ovary | | Testis |
| **STE01** | 32.553/ 89.79/ 56.123 | **STE06** | 43.883/ 90.78/ 57.158 |
| **STE03** | 38.279/ 92.76/ 56.811 | **STE07** | 48.669/ 90.07/ 55.780 |
| **STE05** | 37.663/ 90.94/ 57.531 | **STE08** | 39.004/ 88.68/ 54.893 |
| **Mean** | 37.684/ 91.778/ 56.82 | **STE09** | 43.466/ 88.00/ 55.203 |
| | | **STE10** | 43.148/ 86.83/ 57.063 |
| | | **Mean** | 43.634/ 88.87/ 56.02 |

(C)

| **Number of raw reads [$10^{06}$] / Uniquely mapped reads %/ Reads mapped to genes %** | | | |
|---|---|---|---|
| | Ovary | | Testis |
| **ITE01** | 40.608/ 90.83/ 54.965 | **ITE06** | 38.538/ 88.84/ 57.496 |
| **ITE02** | 34.884/ 93.21/ 56.101 | **ITE07** | 42.537/ 87.94/ 58.489 |
| **ITE03** | 44.289/ 93.30/ 56.780 | **ITE08** | 34.096/ 89.31/ 57.079 |
| **ITE04** | 28.530/ 94.28/ 55.720 | **ITE09** | 24.939/ 87.57/ 58.454 |
| **ITE05** | 70.556/ 93.59/ 56.183 | **ITE10** | 33.684/ 89.69/ 56.984 |
| **Mean** | 43.773/ 93.042/ 55.95 | **Mean** | 32.759/ 88.67/ 57.7 |

**Table S1.** (continued)

(D)

| Number of raw reads [$10^{06}$] / Uniquely mapped reads %/ Reads mapped to genes % | | | |
|---|---|---|---|
| | Ovary | | Testis |
| **HYB1** | 34.826/ 94.04/ 56.104 | **HYB6** | 33.094/ 90.08/ 55.907 |
| **HYB4** | 30.091/ 91.81/ 55.427 | **HYB8** | 35.868/ 87.79/ 50.172 |
| **HYB5** | 29.283/ 93.44/ 56.150 | **HYB10** | 39.132/ 88.05/ 54.278 |
| **HYB7** | 30.696/ 92.07/ 59.013 | **HYB11** | 40.627/ 89.64/ 57.936 |
| **HYB15** | 41.868/ 91.20/ 56.228 | **HYB12** | 40.852/ 89.07/ 58.678 |
| **HYB16** | 78.571/ 92.79/ 57.291 | **Mean** | 40.993/ 89.38/ 55.39 |
| **HYB18** | 69.291/ 93.02/ 57.486 | | |
| **HYB19** | 58.461/ 92.90/ 57.504 | | |
| **Mean** | 50.605/ 92.613/ 56.9 | | |

**Table S2**. Number of annotated genes per chromosome of house sparrow

| Chr | Number of annotated genes | Chr | Number of annotated genes |
|---|---|---|---|
| **1** | 1063 | **17** | 299 |
| **1A** | 774 | **18** | 309 |
| **2** | 1151 | **19** | 305 |
| **3** | 1050 | **20** | 325 |
| **4** | 757 | **21** | 174 |
| **5** | 910 | **22** | 123 |
| **6** | 494 | **23** | 212 |
| **7** | 485 | **24** | 149 |
| **8** | 776 | **25** | 39 |
| **9** | 443 | **26** | 248 |
| **10** | 389 | **27** | 196 |
| **11** | 337 | **28** | 143 |
| **12** | 332 | **Z** | 598 |
| **13** | 339 | **mtDNA** | 11 |
| **14** | 381 | **chrLGE22** | 20 |
| **15** | 361 | | |

**Table S3.** Transcript Integrity Number (TIN).

| Sample | TIN (median) | Sample | TIN (median) |
|---|---|---|---|
| **HTE01** | 82.3965620581 | **HYB7** | 84.9088054893 |
| **HTE02** | 81.0294522839 | **HYB8** | 79.354800142 |
| **HTE03** | 78.2026599633 | **ITE01** | 80.7060705843 |
| **HTE04** | 80.3114393339 | **ITE02** | 81.9869076127 |
| **HTE05** | 78.446957154 | **ITE03** | 80.7380450725 |
| **HTE06** | 79.620645473 | **ITE04** | 77.9153778058 |
| **HTE07** | 76.9344298726 | **ITE05** | 80.0038892674 |
| **HTE08** | 77.5348660828 | **ITE06** | 79.0960303779 |
| **HTE09** | 76.7873621452 | **ITE07** | 78.113096015 |
| **HTE10** | 82.3344852296 | **ITE08** | 78.5822063516 |
| **HYB10** | 76.1242323813 | **ITE09** | 81.3022714567 |
| **HYB11** | 80.2274043433 | **ITE10** | 77.8272960225 |
| **HYB12** | 82.3811990583 | **STE01** | 81.6411126943 |
| **HYB15** | 77.1563412769 | **STE03** | 80.8426466896 |
| **HYB16** | 83.6467004971 | **STE05** | 80.2757441989 |
| **HYB18** | 85.7260398198 | **STE06** | 76.0326224507 |
| **HYB19** | 85.9425671901 | **STE07** | 79.7080461693 |
| **HYB1** | 80.2379060302 | **STE08** | 73.6733563497 |
| **HYB4** | 79.4912608747 | **STE09** | 75.7670969458 |
| **HYB5** | 80.0378109771 | **STE10** | 83.0783909447 |
| **HYB6** | 77.909084622 | | |

**Table S4.** Number of differentially expressed genes and log2 fold change (LFC) with Padj< 0.01 and LFC† > 0.32.

| | Testis | | | Ovary | | |
|---|---|---|---|---|---|---|
| **Comparison** | **Significant** | **LFC > 0** | **LFC < 0** | **Significant** | **LFC > 0** | **LFC < 0** |
| **Spanish - house** | 69 | 45 | 24 | 402 | 311 | 91 |
| **Italian - house** | 1916 | 1092 | 824 | 16 | 6 | 10 |
| **Italian - Spanish** | 2017 | 974 | 1043 | 430 | 65 | 365 |

†LFC: Log Fold Change

**Table S5.** Significant GO categories for the comparison Italian – house testis

| ID | Description | p.adjust | Count |
|---|---|---|---|
| **GO:0019083** | viral transcription | 2.62e-17 | 65 |
| **GO:0006614** | SRP-dependent cotranslational protein targeting to membrane | 2.62e-17 | 62 |
| **GO:0006412** | translation | 8.35e-16 | 109 |
| **GO:0005840** | ribosome | 7.11e-14 | 84 |
| **GO:0044445** | cytosolic part | 1.55e-12 | 46 |
| **GO:0006364** | rRNA processing | 1.39e-09 | 67 |
| **GO:0003735** | structural constituent of ribosome | 2.70e-08 | 153 |
| **GO:0000184** | nuclear-transcribed mRNA catabolic process- nonsense-mediated decay | 3.74e-07 | 37 |
| **GO:0043624** | cellular protein complex disassembly | 4.50e-05 | 25 |
| **GO:0032981** | mitochondrial respiratory chain complex I assembly | 6.23e-05 | 21 |
| **GO:0140053** | mitochondrial gene expression | 0.001405 | 26 |
| **GO:0043488** | regulation of mRNA stability | 0.001 | 32 |
| **GO:0043209** | myelin sheath | 0.003 | 61 |
| **GO:0006401** | RNA catabolic process | 0.004 | 33 |
| **GO:0006119** | oxidative phosphorylation | 0.004 | 16 |
| **GO:0051437** | positive regulation of ubiquitin-protein ligase activity involved in regulation of mitotic cell cycle transition | 0.008 | 23 |
| **GO:0098798** | mitochondrial protein complex | 0.010 | 18 |
| **GO:0002479** | antigen processing and presentation of exogenous peptide antigen via MHC class I- TAP-dependent | 0.010 | 25 |
| **GO:0070069** | cytochrome complex | 0.028 | 9 |
| **GO:0061418** | regulation of transcription from RNA polymerase II promoter in response to hypoxia | 0.031 | 20 |
| **GO:0004040** | amidase activity | 0.037 | 10 |
| **GO:0022900** | electron transport chain | 0.041 | 21 |
| **GO:0033209** | tumor necrosis factor-mediated signaling pathway | 0.041 | 21 |
| **GO:0006754** | ATP biosynthetic process | 0.041 | 12 |
| **GO:1902036** | regulation of hematopoietic stem cell differentiation | 0.041 | 24 |

**Table S5.** (continued)

| | | | |
|---|---|---|---|
| **GO:0032535** | regulation of cellular component size | 0.041 | 13 |
| **GO:0010977** | negative regulation of neuron projection development | 0.042 | 20 |

**Table S6.** Significant GO categories for the comparison Italian – Spanish testis

| ID | Description | p.adjust | Count |
|---|---|---|---|
| **GO:0005840** | ribosome | 6.309e-09 | 78 |
| **GO:0006412** | translation | 2.304e-06 | 92 |
| **GO:0019083** | viral transcription | 7.5207e-06 | 50 |
| **GO:0043624** | cellular protein complex disassembly | 7.520e-06 | 27 |
| **GO:0044445** | cytosolic part | 1.172e-05 | 38 |
| **GO:0006614** | SRP-dependent cotranslational protein targeting to membrane | 3.745e-05 | 46 |
| **GO:0003735** | structural constituent of ribosome | 4.463e-05 | 144 |
| **GO:0043209** | myelin sheath | 0.0002 | 65 |
| **GO:0010977** | negative regulation of neuron projection development | 0.0092 | 22 |
| **GO:0006364** | rRNA processing | 0.0092 | 53 |
| **GO:0032981** | mitochondrial respiratory chain complex I assembly | 0.0092 | 18 |
| **GO:0140053** | mitochondrial gene expression | 0.011 | 25 |
| **GO:0000027** | ribosomal large subunit assembly | 0.025 | 11 |
| **GO:0000184** | nuclear-transcribed mRNA catabolic process- nonsense-mediated decay | 0.029 | 29 |
| **GO:0006754** | ATP biosynthetic process | 0.033 | 13 |

**Table S7.** Significant GO categories for the comparison Italian – house ovary

| ID | Description | p.adjust | Count |
|---|---|---|---|
| **GO:0015081** | sodium ion transmembrane transporter activity | 0.033 | 2 |
| **GO:0086010** | membrane depolarization during action potential | 0.033 | 2 |
| **GO:0005261** | cation channel activity | 0.033 | 2 |
| **GO:0005244** | voltage-gated ion channel activity | 0.047 | 2 |
| **GO:0008217** | regulation of blood pressure | 0.047 | 2 |
| **GO:0098660** | inorganic ion transmembrane transport | 0.047 | 2 |

**Table S8.** Significant GO categories for the comparison Italian – Spanish ovary

| ID | Description | p.adjust | Count |
|---|---|---|---|
| **GO:0001933** | negative regulation of protein phosphorylation | 0.048 | 19 |

**Table S9.** Number and percentage of genes in each inheritance category for differentially expressed genes with Padj < 0.01.

| | **Testis** | **Ovary** |
|---|---|---|
| **Inheritance** | **Italian** | **Italian** |
| **Conserved** | 10237 (89.09%) | 11596 (98.25%) |
| **Additive** | 1 (0.009%) | 8 (0.07%) |
| **house dominant** | 35 (0.30%) | 198 (1.68%) |
| **Spanish dominant** | 12 (0.10%) | 3 (0.025%) |
| **Underdominant** | 582 (5.06%) | 2 (0.017%) |
| **Overdominant** | 626 (5.45%) | 0 (0%) |

**Table S10.** Significant GO categories for over-dominant genes in testis of Italian sparrow

| ID | Description | p.adjust | Count |
|---|---|---|---|
| **GO:0005840** | ribosome | 1.123e-20 | 60 |
| **GO:0006412** | translation | 1.123e-20 | 71 |
| **GO:0006614** | SRP-dependent cotranslational protein targeting to membrane | 1.238-17 | 42 |
| **GO:0019083** | viral transcription | 2.089-15 | 41 |
| **GO:0044445** | cytosolic part | 1.179-14 | 33 |
| **GO:0003735** | structural constituent of ribosome | 1.437-14 | 91 |
| **GO:0006364** | rRNA processing | 2.999e-11 | 42 |
| **GO:0043624** | cellular protein complex disassembly | 4.186e-11 | 22 |
| **GO:0140053** | mitochondrial gene expression | 4.648e-10 | 23 |
| **GO:0043209** | myelin sheath | 9.759e-09 | 42 |
| **GO:0032981** | mitochondrial respiratory chain complex I assembly | 1.202e-08 | 17 |
| **GO:0000184** | nuclear-transcribed mRNA catabolic process- nonsense-mediated decay | 6.760e-08 | 24 |
| **GO:0000027** | ribosomal large subunit assembly | 1.751e-05 | 10 |
| **GO:0005743** | mitochondrial inner membrane | 1.974e-05 | 24 |
| **GO:0098798** | mitochondrial protein complex | 3.091e-05 | 14 |
| **GO:0005759** | mitochondrial matrix | 3.091e-05 | 19 |
| **GO:0022900** | electron transport chain | 7.211e-05 | 16 |
| **GO:0006119** | oxidative phosphorylation | 8.833e-05 | 12 |
| **GO:0070069** | cytochrome complex | 0.0001 | 8 |
| **GO:0006457** | protein folding | 0.0001 | 30 |
| **GO:0006754** | ATP biosynthetic process | 0.0002 | 10 |
| **GO:0031012** | extracellular matrix | 0.0004 | 38 |
| **GO:1990204** | oxidoreductase complex | 0.0005 | 16 |
| **GO:0002479** | antigen processing and presentation of exogenous peptide antigen via MHC class I- TAP-dependent | 0.0006 | 16 |
| **GO:0015078** | hydrogen ion transmembrane transporter activity | 0.001 | 20 |
| **GO:0004040** | amidase activity | 0.001 | 8 |
| **GO:0051437** | positive regulation of ubiquitin-protein ligase activity involved in regulation of mitotic cell cycle transition | 0.002 | 14 |

**Table S10.** (continued)

| | | | |
|---|---|---|---|
| **GO:0007339** | binding of sperm to zona pellucida | 0.002 | 10 |
| **GO:0061418** | regulation of transcription from RNA polymerase II promoter in response to hypoxia | 0.002 | 13 |
| **GO:0043488** | regulation of mRNA stability | 0.004 | 17 |
| **GO:0006521** | regulation of cellular amino acid metabolic process | 0.005 | 13 |
| **GO:0033209** | tumor necrosis factor-mediated signaling pathway | 0.005 | 13 |
| **GO:0006413** | translational initiation | 0.005 | 20 |
| **GO:0005844** | polysome | 0.005 | 12 |
| **GO:0002181** | cytoplasmic translation | 0.005 | 9 |
| **GO:0042787** | protein ubiquitination involved in ubiquitin-dependent protein catabolic process | 0.008 | 10 |

**Table S11.** Number and percentage of genes in each inheritance category in experimental F1 hybrids.

| | **Testis** | **Ovary** |
|---|---|---|
| **Inheritance** | **EXPERIMENTAL** | **EXPERIMENTAL** |
| **Conserved** | 11515 (99.14%) | 11600 (96.9%) |
| **Additive** | 6 (0.05%) | 272 (2.27%) |
| **house dominant** | 45 (0.39%) | 94 (0.78%) |
| **Spain dominant** | 4 (0.03%) | 6 (0.05%) |
| **Underdominant** | 12 (0.1%) | 2 (0.17%) |
| **Overdominant** | 33 (0.27%) | 2 (0.17%) |

**Table S12.** Significant GO categories for the comparison of experimental F1 – house testis

| ID | Description | p.adjust | Count |
|---|---|---|---|
| **GO:0005578** | proteinaceous extracellular matrix | 4.451e-07 | 11 |
| **GO:0005201** | extracellular matrix structural constituent | 2.938e-06 | 10 |
| **GO:1903561** | extracellular vesicle | 1.022e-05 | 10 |
| **GO:0005581** | collagen trimer | 1.056e-05 | 9 |
| **GO:0030574** | collagen catabolic process | 1.149e-05 | 6 |
| **GO:0031012** | extracellular matrix | 3.226e-05 | 11 |
| **GO:0044420** | extracellular matrix component | 4.494e-05 | 5 |
| **GO:0005518** | collagen binding | 7.980e-05 | 6 |
| **GO:0005509** | calcium ion binding | 8.882e-05 | 16 |
| **GO:0009612** | response to mechanical stimulus | 0.0008 | 6 |
| **GO:0001501** | skeletal system development | 0.0020 | 7 |
| **GO:0042060** | wound healing | 0.0024 | 8 |
| **GO:0009653** | anatomical structure morphogenesis | 0.0025 | 14 |
| **GO:0008201** | heparin binding | 0.0045 | 6 |
| **GO:0001558** | regulation of cell growth | 0.0045 | 6 |
| **GO:0005520** | insulin-like growth factor binding | 0.0067 | 3 |
| **GO:0040011** | locomotion | 0.0074 | 11 |
| **GO:0001568** | blood vessel development | 0.0081 | 5 |
| **GO:0045089** | positive regulation of innate immune response | 0.0085 | 4 |
| **GO:0001666** | response to hypoxia | 0.0093 | 7 |
| **GO:0065008** | regulation of biological quality | 0.0104 | 10 |
| **GO:0001968** | fibronectin binding | 0.0106 | 3 |
| **GO:0051674** | localization of cell | 0.0121 | 8 |
| **GO:0031295** | T cell costimulation | 0.0121 | 3 |
| **GO:0007268** | chemical synaptic transmission | 0.01669 | 5 |
| **GO:0009888** | tissue development | 0.01992 | 7 |
| **GO:0016525** | negative regulation of angiogenesis | 0.02163 | 4 |
| **GO:0001775** | cell activation | 0.0243 | 5 |
| **GO:0052548** | regulation of endopeptidase activity | 0.0243 | 3 |

**Table S12.** (continued)

| | | | |
|---|---|---|---|
| **GO:0007409** | axonogenesis | 0.0252 | 4 |
| **GO:0045471** | response to ethanol | 0.0263 | 5 |
| **GO:0050840** | extracellular matrix binding | 0.0294 | 3 |
| **GO:0006955** | immune response | 0.0316 | 7 |
| **GO:0005254** | chloride channel activity | 0.03166 | 3 |
| **GO:0007568** | aging | 0.033 | 7 |
| **GO:0010811** | positive regulation of cell-substrate adhesion | 0.033 | 3 |
| **GO:0031099** | regeneration | 0.035 | 5 |
| **GO:0008076** | voltage-gated potassium channel complex | 0.035 | 3 |
| **GO:0071805** | potassium ion transmembrane transport | 0.035 | 4 |
| **GO:0048513** | animal organ development | 0.039 | 8 |
| **GO:0005788** | endoplasmic reticulum lumen | 0.041 | 4 |
| **GO:0032501** | multicellular organismal process | 0.043 | 12 |
| **GO:0050900** | leukocyte migration | 0.044 | 3 |
| **GO:0007160** | cell-matrix adhesion | 0.046 | 4 |
| **GO:0007566** | embryo implantation | 0.046 | 3 |
| **GO:0030276** | clathrin binding | 0.046 | 3 |

**Table S13.** Significant GO categories for the comparison of experimental F1 – Spanish testis

| ID | Description | p.adjust | Count |
|---|---|---|---|
| **GO:0030574** | collagen catabolic process | 0.010 | 6 |
| **GO:0042060** | wound healing | 0.03 | 12 |
| **GO:0008277** | regulation of G-protein coupled receptor protein signaling pathway | 0.043 | 5 |
| **GO:0005201** | extracellular matrix structural constituent | 0.047 | 9 |
| **GO:0031099** | regeneration | 0.047 | 9 |
| **GO:0001568** | blood vessel development | 0.047 | 7 |
| **GO:0070374** | positive regulation of ERK1 and ERK2 cascade | 0.047 | 7 |

**Table S14.** Significant GO categories for the comparison experimental F1 – Spanish ovary

| ID | Description | p.adjust | Count |
|---|---|---|---|
| **GO:0033280** | response to vitamin D | 0.02 | 4 |
| **GO:1903508** | positive regulation of nucleic acid-templated transcription | 0.02 | 15 |
| **GO:0010629** | negative regulation of gene expression | 0.02 | 17 |
| **GO:0051216** | cartilage development | 0.031 | 5 |
| **GO:0042981** | regulation of apoptotic process | 0.031 | 11 |
| **GO:1902911** | protein kinase complex | 0.032 | 3 |
| **GO:0032410** | negative regulation of transporter activity | 0.032 | 3 |
| **GO:0045429** | positive regulation of nitric oxide biosynthetic process | 0.032 | 3 |
| **GO:0048856** | anatomical structure development | 0.032 | 11 |
| **GO:0033993** | response to lipid | 0.032 | 4 |
| **GO:0004222** | metalloendopeptidase activity | 0.032 | 6 |
| **GO:0005520** | insulin-like growth factor binding | 0.032 | 3 |
| **GO:0015807** | L-amino acid transport | 0.032 | 3 |
| **GO:0045840** | positive regulation of mitotic nuclear division | 0.032 | 3 |
| **GO:0033574** | response to testosterone | 0.032 | 4 |
| **GO:0042127** | regulation of cell proliferation | 0.032 | 9 |
| **GO:0001067** | regulatory region nucleic acid binding | 0.032 | 10 |
| **GO:0010042** | response to manganese ion | 0.032 | 3 |
| **GO:0035116** | embryonic hindlimb morphogenesis | 0.032 | 3 |
| **GO:0005581** | collagen trimer | 0.04 | 6 |
| **GO:0002684** | positive regulation of immune system process | 0.043 | 6 |
| **GO:0004252** | serine-type endopeptidase activity | 0.043 | 6 |
| **GO:0005125** | cytokine activity | 0.0443 | 4 |
| **GO:1903507** | negative regulation of nucleic acid-templated transcription | 0.0443 | 11 |
| **GO:0001503** | ossification | 0.0453 | 6 |

**Table S14.** (continued)

| | | | |
|---|---|---|---|
| **GO:2000113** | negative regulation of cellular macromolecule biosynthetic process | 0.0453 | 11 |
| **GO:0032148** | activation of protein kinase B activity | 0.046 | 3 |
| **GO:0050829** | defense response to Gram-negative bacterium | 0.046 | 3 |

**Table S15.** Significant GO categories for over-dominant genes in testis of experimental F1 hybrid

| ID | Description | p.adjust | Count |
|---|---|---|---|
| **GO:0030574** | collagen catabolic process | 0.0002 | 4 |
| **GO:0042060** | wound healing | 0.0004 | 6 |
| **GO:0005520** | insulin-like growth factor binding | 0.0004 | 3 |
| **GO:0005578** | proteinaceous extracellular matrix | 0.0004 | 5 |
| **GO:0001501** | skeletal system development | 0.0005 | 5 |
| **GO:0044420** | extracellular matrix component | 0.001 | 3 |
| **GO:0001568** | blood vessel development | 0.001 | 4 |
| **GO:0048589** | developmental growth | 0.001 | 5 |
| **GO:0001666** | response to hypoxia | 0.001 | 5 |
| **GO:0050840** | extracellular matrix binding | 0.001 | 3 |
| **GO:0001558** | regulation of cell growth | 0.002 | 4 |
| **GO:0005581** | collagen trimer | 0.002 | 4 |
| **GO:0009888** | tissue development | 0.002 | 5 |
| **GO:0005201** | extracellular matrix structural constituent | 0.003 | 4 |
| **GO:0045597** | positive regulation of cell differentiation | 0.006 | 3 |
| **GO:0016525** | negative regulation of angiogenesis | 0.007 | 3 |
| **GO:0061448** | connective tissue development | 0.013 | 3 |
| **GO:0005788** | endoplasmic reticulum lumen | 0.013 | 3 |
| **GO:0031594** | neuromuscular junction | 0.013 | 3 |

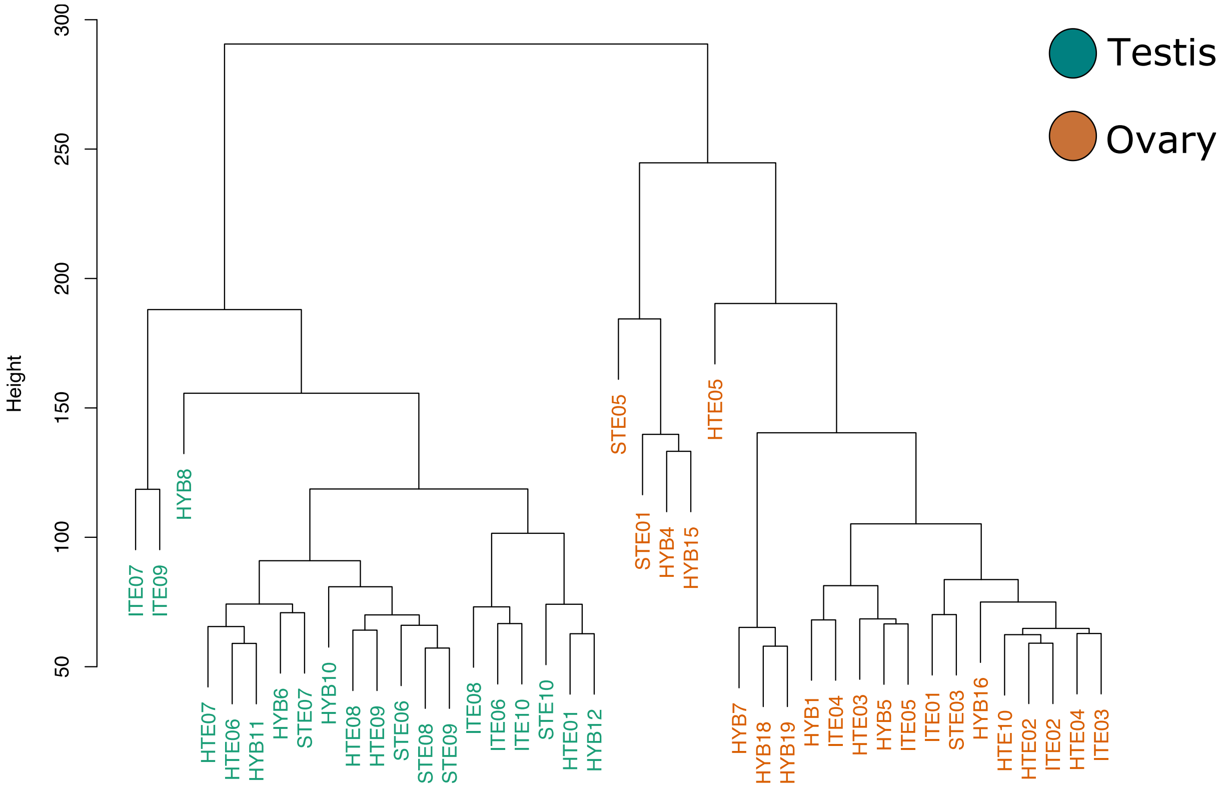


**Figure S1.** Cluster dendrogram of sample clustering separates the samples into two clusters for testis and ovary.

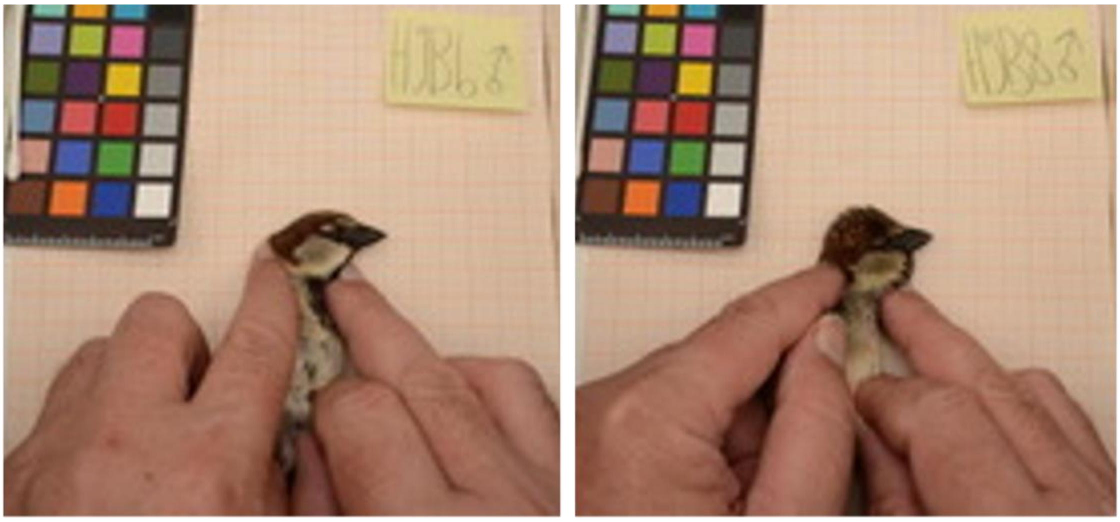

**Figure S2.** Pictures of two of the experimental F1 hybrid males (HYB6 and HYB8).

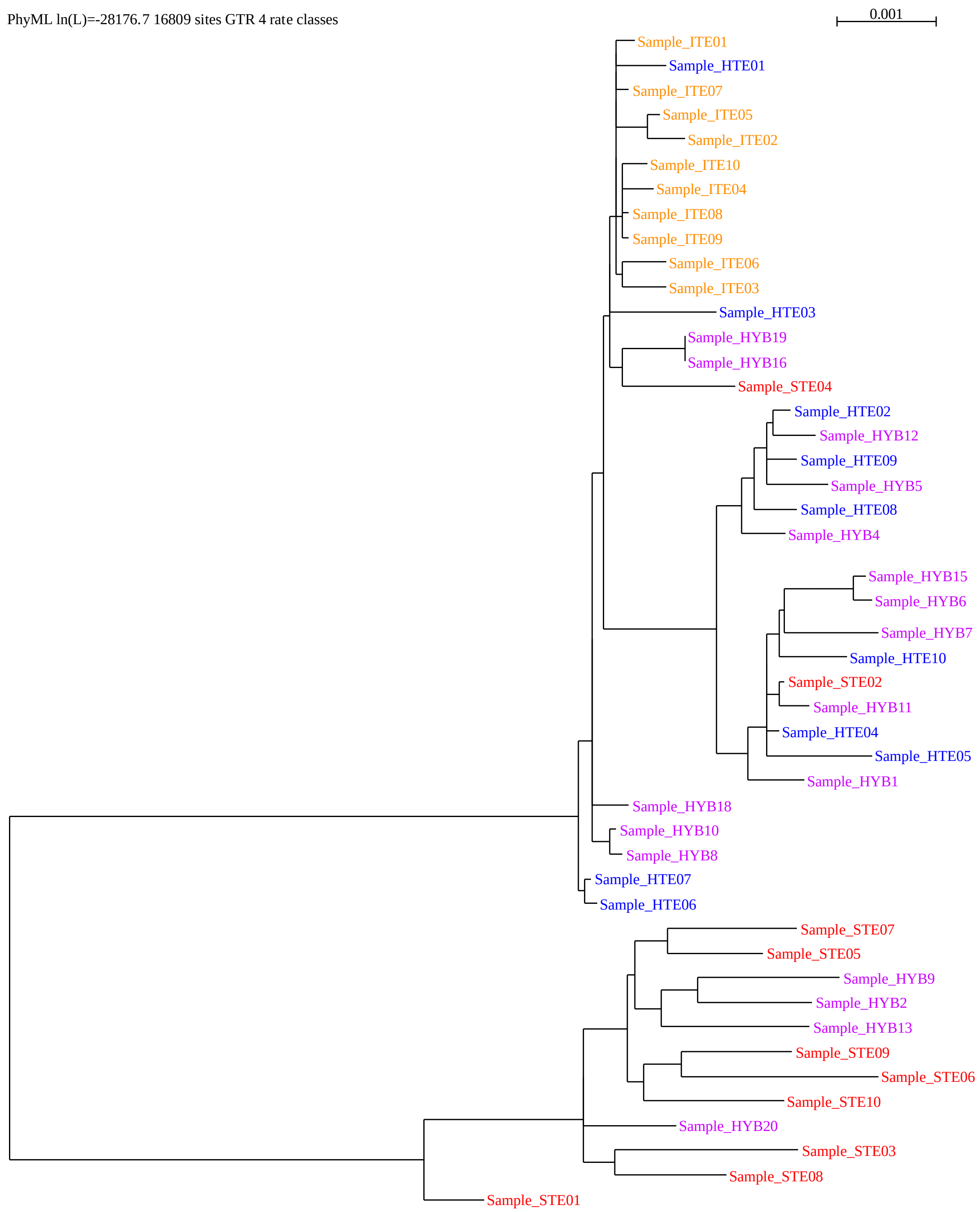


**Figure S3**. Mitochondrial tree of all samples in this study. Four of the experimental F1 samples (HYB9, HYB2, HYB13 and HYB20) cluster with Spanish sparrows and two Spanish sparrow samples (STE02 and STE04) cluster with house mtDNA. These samples were excluded from subsequent analyses.

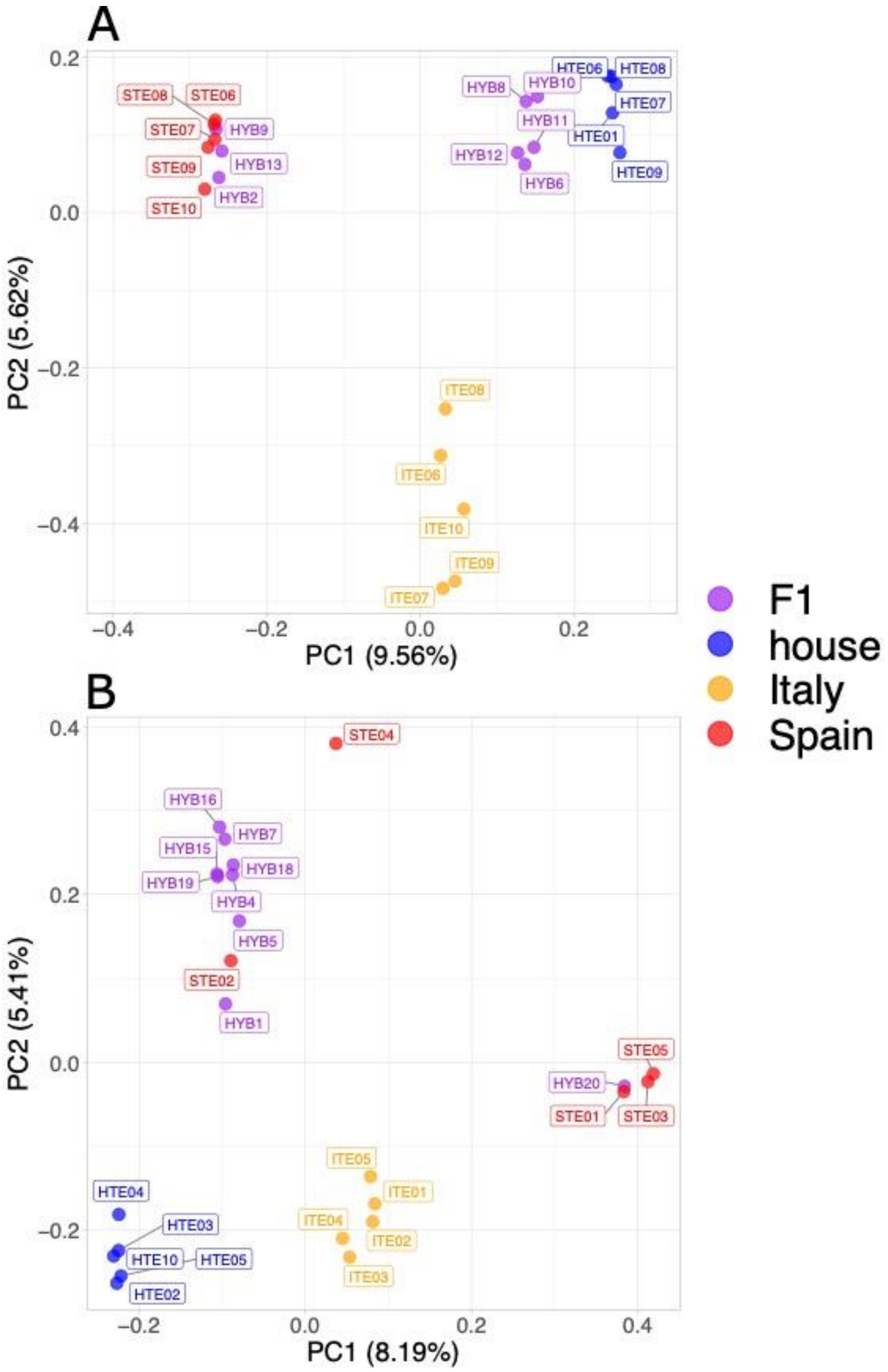


**Figure S4.** Principal component analysis of autosomal SNPs. A) Testis B) Ovary

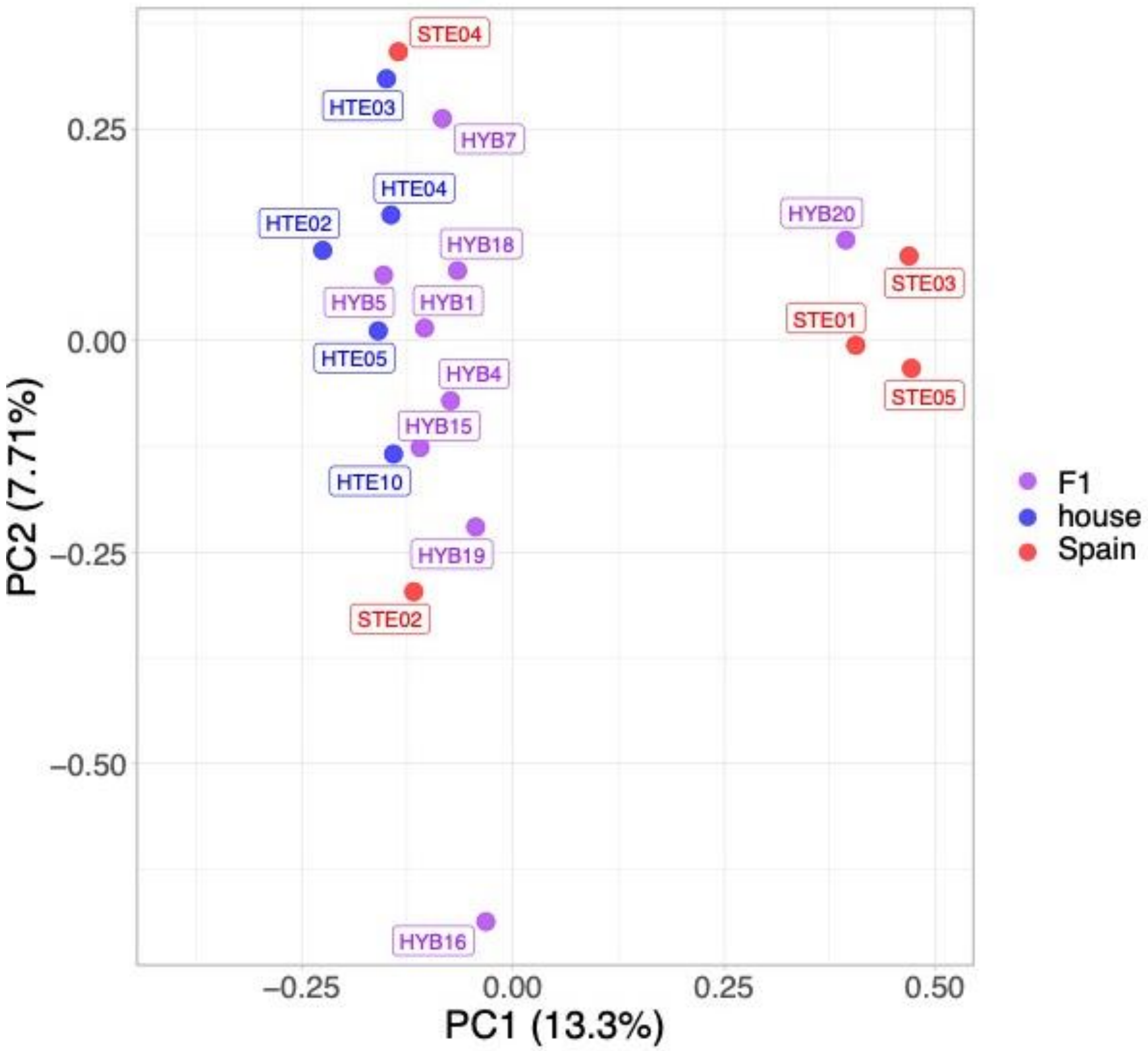


**Figure S5**. PCA of the ovary Z chromosome.

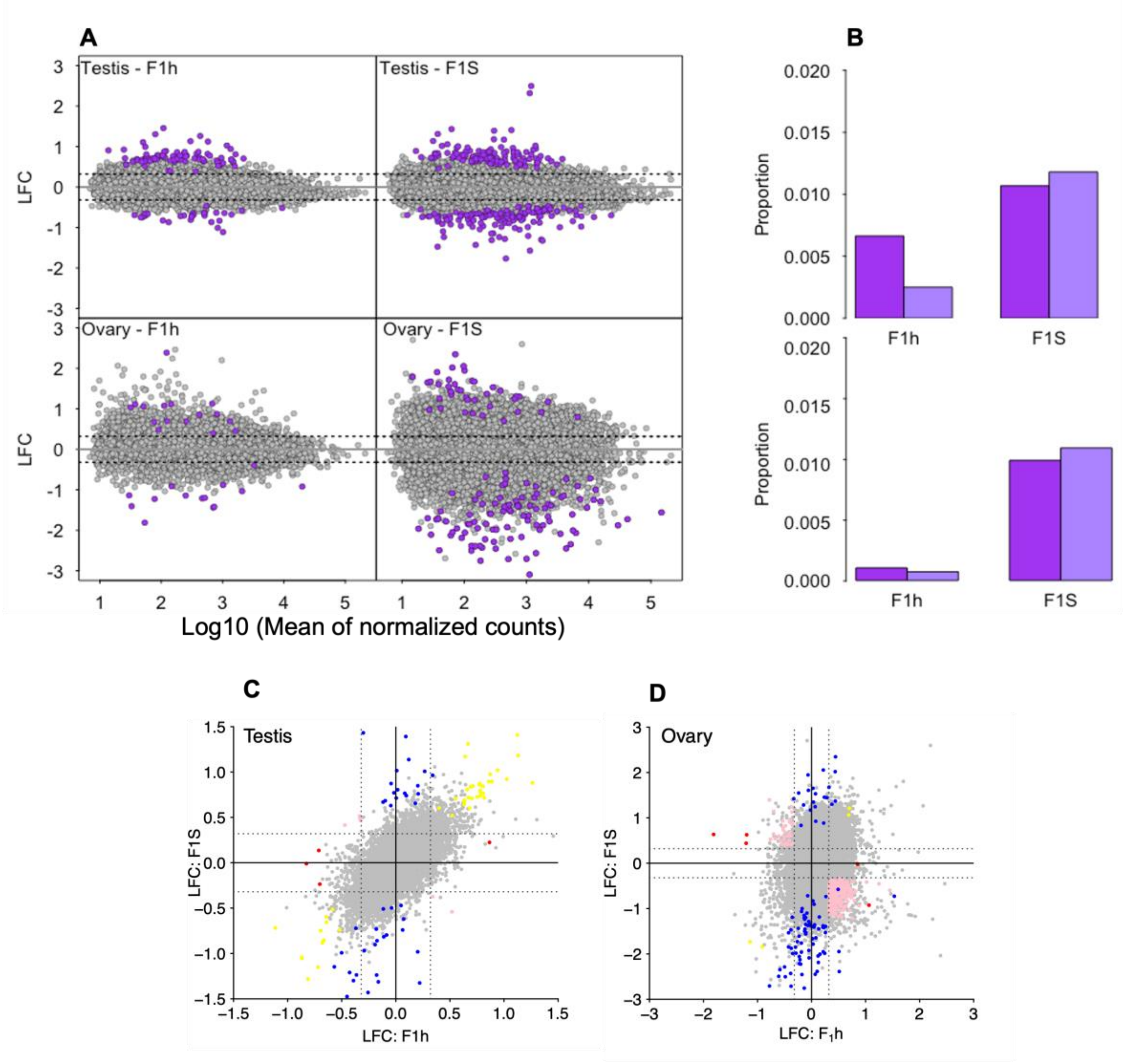


**Figure S6**. Gene expression and inheritance pattern in experimental F1 hybrids in comparison to parental species, house (F1h) and Spanish (F1S) sparrows. A) Log2 fold change (LFC) as a function of the mean of normalized counts. B) Proportion of up- (dark purple) and down- (light purple) regulated genes. C) Inheritance pattern of experimental F1 hybrids for testis and D) for ovary. Grey points in each graph depicts the total number of genes studied for gene expression with those colored representing the ones significantly different from parental species to be divided into each of the inheritance categories (Conserved: grey, additive: pink, house dominant: blue, Spain dominant: red, transgressive (over-dominant and under-dominant: yellow). Grey dotted lines indicate the log2 fold change of 0.32 used for classification of inheritance pattern.
